# Membranous interacting partners of phage-type plastid RNA polymerase have limited impact on plastid gene expression during chloroplast development

**DOI:** 10.1101/2025.05.13.653719

**Authors:** Yushi Kurotaki, Atsuo S. Nishino, Sho Fujii

## Abstract

In vascular plants, genes in the plastid genome are transcribed by two types of RNA polymerases, namely bacterial-type nuclear-encoded and phage-type plastid-encoded plastid RNA polymerases (NEP and PEP, respectively). Eudicots, including Arabidopsis, carry two isoforms of NEP, RPOTp and RPOTmp. NEPs transcribe multiple plastid-encoded genes including subunits of PEP and translocon, thus are indispensable for the maintenance of plastids. However, regulatory mechanisms of NEPs are largely unknown. RPOTmp transcribes the 16S rRNA gene from a specific promoter in the seeds during vernalization, and its mutation in Arabidopsis retards the chloroplast development. As interacting partners of RPOTmp, two NEP-INTERACTING PROTEINs (NIP1 and NIP2) have been identified and suggested to suppress RPOTmp activity by tethering RPOTmp to the thylakoid membrane during chloroplast development in the presence of light, but their precise roles in transcriptional regulation remain to be addressed. From these previous reports, we hypothesize that functions of RPOTmp would depend on the light conditions and expression of NIPs. To gain insight into how RPOTmp is controlled, we performed functional analysis of RPOTmp and NIPs using Arabidopsis mutants during germination in the dark and de-etiolation processes under light. We found that RPOTmp-dependent transcription of 16S rRNA is active in imbibed seeds and remains basal level throughout the post-germination processes, regardless of light conditions. We also demonstrated a limited impact of NIPs on RPOTmp function during these processes. Our phylogenetic analysis indicates that NIPs have distinct evolutionary profiles compared to RPOTmp, and Arabidopsis is unlikely to have additional NIP-like proteins in plastids. Based on these findings, we propose a modified model of RPOTmp regulation during chloroplast development: RPOTmp activity remains stable throughout the process of the chloroplast differentiation and is unaffected by light and NIPs.

## Introduction

Plastid genome encodes multiple genes for photosynthesis and its own gene expression system. In seed plants, these genes are transcribed by two types of RNA polymerase, namely bacterial type plastid-encoded plastid RNA polymerase (PEP) and bacteriophage-type nuclear-encoded plastid RNA polymerase (NEP) (Börner et al. 2015). PEP is responsible for transcription of photosynthesis-associated genes and some tRNA genes and its promoter recognition depends on sigma factors (Shiina et al. 2009; Palomar et al. 2022). In angiosperms, ∼15 peripheral subunits are identified as essential factors for PEP activation in chloroplasts (Pfalz et al. 2006; Pfalz and Pfannschmidt 2013; Wu et al. 2024; Vergara-Cruces et al. 2024; Prado et al. 2024; Wang et al. 2024). In contrast to PEP, NEP transcribes a broader range of plastid-encoded genes involved in photosynthesis, gene expression, protein import, and other metabolic processes (Börner et al. 2015). NEP functions as a single-subunit protein by recognizing at least three types of promoters, namely type Ia (YRTa), type Ib (GGA + YRTa), and type II (GGA) (Swiatecka-Hagenbruch et al. 2007). In Arabidopsis, two isoforms of NEP are targeted to plastids. Arabidopsis mutant lacking two NEP isoforms shows seedling-lethal phenotype (Hricová et al. 2006), suggesting crucial roles of NEP in plastid functionality and plant development.

NEP proteins in plastids share evolutionary ancestry with mitochondrial RNA polymerase. In prasinophytes and green algae, a single gene of phage-type RNA polymerase (RPOT) is identified and is considered to be solely targeted to mitochondria (Yin et al. 2010; Liere et al. 2011; Richter et al. 2014). By contrast, two clades of RPOT are found in Lycophytes and Spermatophyta (Yin et al. 2010; Richter et al. 2014). One is called RPOTp and localized in plastids. The other consists of mitochondria- and dual-targeted proteins, which are called RPOTm and RPOTmp, respectively (Hedtke et al. 1997, 2000). In contrast to widely conserved RPOTm, RPOTmp are specific to eudicots but not found in monocots or Lycophytes (Yin et al. 2010; Richter et al. 2014), thus it is likely that plastid in eudicots acquired additional isoforms of RPOT in a later step of the evolution process. These insights indicate that the addition of plastid-targeted RPOT may contribute to the adaptation of photosynthetic organisms in various environmental conditions.

In Arabidopsis, RPOTp-less mutant (*rpotp*) shows pale-green phenotype and decreased expression level of multiple plastid-encoded genes in the beginning phase of germination (Hricová et al. 2006), probably due to the low transcriptional levels of housekeeping genes. Suppression of these genes becomes milder in a later developmental stage. Given that the phenotype of the *rpotp* knockout mutant is not as severe as the seedling-leathal *rpotp rpotmp* double mutant (Hricová et al. 2006), RPOTmp may contribute to maintaining the minimal function of plastids in the *rpotp* mutant. In RPOTmp-deficient mutant (*rpotmp*), root elongation is significantly repressed and the greening of etiolated seedlings after light illumination is also retarded, although the photosynthetic function is recovered in the later developmental steps (Baba et al. 2004). RPOTmp is considered to play an important role in mitochondria, because the *rpotmp* mutation strongly disturbs mitochondrial gene expression pattern (Kühn et al. 2007) and the introduction of mitochondrial-targeted RPOTmp complements the growth phenotype of *rpotmp* mutant (Tarasenko et al. 2016). Transcription initiation of RPOTmp in vitro is less sequence-specific than that of other RPOTs (Bohne et al. 2016), but in plastids, RPOTmp actively transcribes 16S rRNA gene (*rrn16*) from a specific promoter (PC promoter) (Courtois et al. 2007). Also, RPOTmp potentially transcribes multiple housekeeping genes in plastids, such as *accD, clpP*, and *rpoB* (Gorbenko et al. 2024). Expression of RpoB, a core subunit of PEP, fully relies on RPOTp and/or RPOTmp (Zhelyazkova et al. 2012), thus NEPs are critical for activate entire gene expression system in plastids.

Despite the importance of NEP in plastids, little is known about the functional regulation of these proteins. Courtois et al. (2007) demonstrated that RPOTmp-dependent transcription of *rrn16* from PC promoter is activated during seed imbibition, and this activity is downregulated within 2 days of germination. By contrast, the RPOTmp protein level is increased in this stage of germination (Demarsy et al. 2006), suggesting an existence of transcriptional regulators which suppress RPOTmp activity.

In spinach, membrane association of RPOTmp proteins is weak in etioplasts and strong in chloroplasts (Azevedo et al. 2008), thus it may be possible that the light-dependent changes of intraorganellar localization contribute to the regulation of this RNA polymerase. Membranous RING-H2 proteins named NEP-INTERACTING PROTEINs (NIPs) have been isolated as interacting partners of RPOTmp in Arabidopsis and spinach (Azevedo et al. 2008). NIP has three transmembrane domains in its N-terminus and is considered to interact with RPOTmp via its C-terminal RING domain. Accumulation levels of NIP protein anticorrelate with the pattern of RPOTmp-dependent transcription of *rrn16* during the development of chloroplasts. Based on these findings, it has been hypothesized that light triggers NIP-mediated RPOTmp recruitment to the thylakoid membrane and suppresses the transcriptional activity of RPOTmp in the initial stages of chloroplast differentiation (Azevedo et al. 2008), although the direct evidence to evaluate this hypothesis remains missing.

To understand the mechanism of NEP regulation, we aimed to address the contribution of NIP proteins in RPOTmp functionality during chloroplast development in this study. Since Arabidopsis carries two paralogs of NIP, namely NIP1 and NIP2, we generated double mutant (*nip1 nip2*) of two NIP genes to clarify the physiological impact of these thylakoid RING proteins. To elucidate whether RPOTmp and NIP contributes the germination process or chloroplast differentiation, we investigated the transcriptional profile of plastid-encoded genes in *nip1 nip2* in the process of germination, etiolated growth in darkness, and etioplast-to-chloroplast differentiation under light exposure. Our results demonstrated that RPOTmp shows high activity in imbibed seeds and functions constantly throughout the etioplast formation and light-dependent chloroplast differentiation. We did not find any significant influence of NIP proteins on RPOTmp functions during these processes. Also, our molecular phylogenetic analysis indicated that gene orthologs of *NIP1* and *NIP2* are widely conserved in angiosperms including monocots, which do not have RPOTmp. Taken together, we propose that NIP proteins might be acquired before the emergence of RPOTmp, and may play limited roles on functional regulation of RPOTmp during the chloroplast development.

## Results

### *NIP1* and *NIP2* are differentially regulated in response to light

Previous study showed that accumulation of NIP proteins is induced during chloroplast development (Azevedo et al. 2008), but it remains unclear which isoform of NIP genes is responsible for the increase of NIP protein levels. To clarify the expression regulation of *NIP1* and *NIP2* genes during chloroplast development, we evaluated the transcript levels of these genes in wild-type seedlings during the process of etiolation and deetiolation. Given the high similarity of coding sequence of two NIP genes, we targeted 5’-UTR of NIP genes for RT-qPCR analysis (Fig. 1A). In darkness, the transcript levels of both *NIP1* and *NIP2* were mostly stable (Fig. 1B). *NIP1* mRNA level was increased twice at the onset of light illumination, whereas the *NIP2* mRNA level was gradually decreased after light exposure, indicating that the expression of two NIP isoforms is differentially regulated in response to light. *NIP1* is likely to be responsible for NIP protein accumulation during chloroplast development.

**Figure 1.**
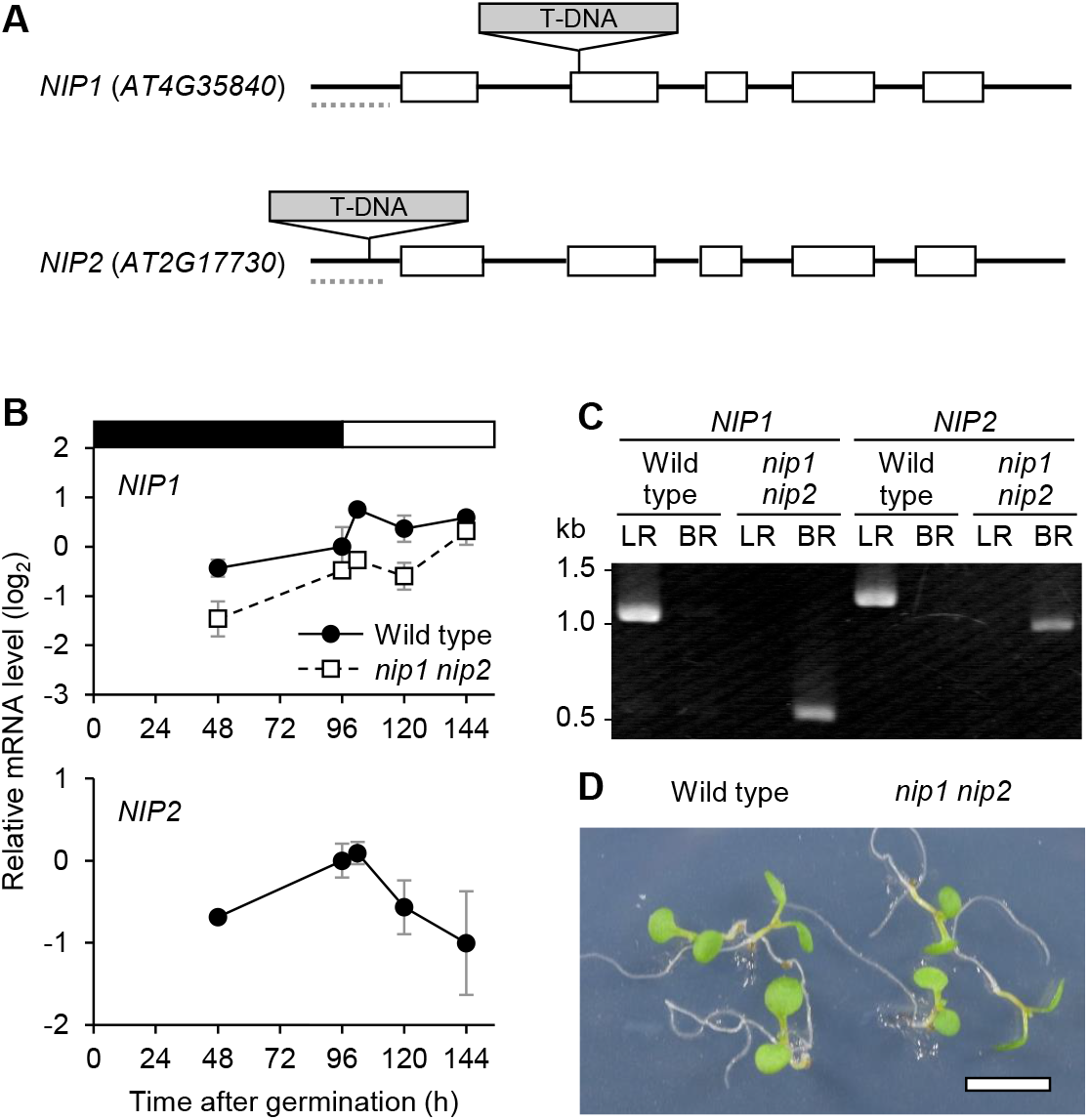
Characterization of *NIP1* and *NIP2* genes and their mutants in Arabidopsis. (A) Schematic diagram of *NIP1* and *NIP2* genes. Open boxes and black lines indicate exons and UTRs or introns, respectively. Gray boxes with triangles show the T-DNA insertion sites in *nip1* (SALK_057350) and *nip2* (SALK_137112) mutants. Broken gray lines indicate target regions of RT-qPCR analysis. (B) RT-qPCR analysis of *NIP1* and *NIP2* in wild-type and *nip1 nip2* seedlings. Seedlings were incubated in darkness for 96 h and then illuminated under continuous light for indicated time length. Transcript levels were normalized to the averaged level of *ACTIN8* and *ELONGATION FACTOR 1α* and presented as the difference from the wild type at 96 h of dark growth. Data are mean ± SE from three biological replicates. (C) Genotyping PCR to confirm T-DNA insertions in *NIP1* and *NIP2* genes. Wild-type and *nip1 nip2* seedlings were utilized for analysis. LR and BR indicate primer sets used in PCR to detect NIP gene without or with T-DNA insertion, respectively. (D) Phenotype of *nip1 nip2* mutant seedlings grown for 4 d under continuous white light. Bar = 5 mm.

### Loss of both NIP1 and NIP2 have invisible impact on cotyledon greening

To assess the roles of NIP proteins in plastids, we established double mutant lacking NIP1 and NIP2 (*nip1 nip2*) by crossing T-DNA insertion mutants of NIP1 (*nip1*) and NIP2 (*nip2*) (Fig. 1A). Our PCR analysis confirmed the insertion of T-DNA fragments in the second exon of *NIP1* and the 5’-UTR of *NIP2* (Fig 1C). To evaluate the expression level of NIP genes in *nip1 nip2* mutant, we performed RT-qPCR analysis of *NIP1* and *NIP2* (Fig 1B). The transcript level of *NIP2* was below detection threshold in the double mutant. *NIP1* mRNA content in *nip1 nip2* was not largely decreased from the wild-type level, probably because our target of RT-qPCR was the upstream of T-DNA insertion site. Considering that the T-DNA insertion site of *nip1* mutant is located at the second exon, we concluded that *nip1 nip2* double mutant does not express functional NIP proteins. Under continuous light conditions, the *nip1 nip2* double mutant did not show obvious phenotype (Fig 1D), indicating that the greening of cotyledons was unaffected by the complete loss of two NIP proteins.

### NIP proteins have limited impact on accumulation of RPOTmp-dependent transcripts

RPOTmp transcribes *rrn16* gene from a specific promoter called PC promoter (Courtois et al. 2007). The transcriptional activity of RPOTmp on the PC promoter is particularly strong at the initial stage of germination and decreased within 2 days in Arabidopsis (Courtois et al. 2007; Azevedo et al. 2008). To quantify RPOTmp-dependent transcripts in vivo by using RT-qPCR, the level of transcript initiated from upstream promoters must be subtracted from the abundance of the transcript detected by PC promoter-specific primers (Tarasenko et al. 2016). We confirmed that the abundance of PC-promoter-initiated transcripts was larger in wild type compared to the RPOTmp-deficient mutant (*rpotmp*) throughout the process of etiolation and deetiolation (Fig. 2A). In wild type, the transcript levels of the PC-promoter region were particularly elevated in vernalized seeds consistent to the previous studies (Courtois et al. 2007; Azevedo et al. 2008). This result demonstrated that RPOTmp-dependent transcription of *rrn16* was unaffected by light treatment on etiolated seedlings.

**Figure 2.**
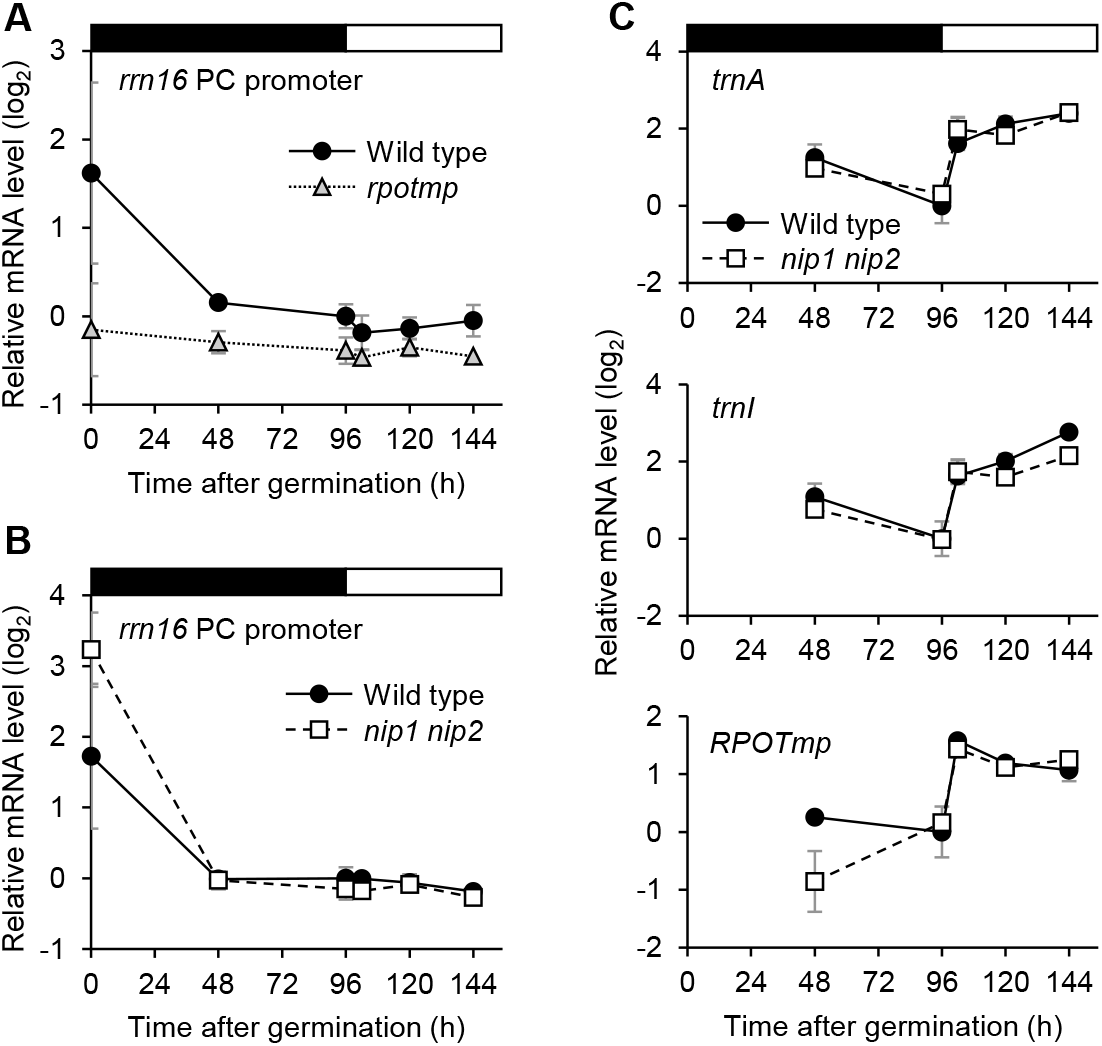
Effect of NIPs on RPOTmp-dependent gene expression. (A) RT-qPCR analysis of PC-promoter initiated transcripts in the wild type and *rpotmp* mutant. (B) RT-qPCR analysis of PC-promoter initiated transcripts in the wild type and *nip1 nip2* mutant. In A and B, levels of PC promoter-initiated transcript were normalized to the background level of *rrn16* transcript as described (Tarasenko et al. 2016) and presented as the difference from the wild type at 96 h of dark growth. 0 h indicates imbibed seeds before incubation at 23°C. (C) RT-qPCR analysis of plastid-encoded tRNAs and *RPOTmp*. Transcript levels were normalized to the averaged level of *ACTIN8* and *ELONGATION FACTOR 1α* and presented as the difference from the wild type at 96 h of dark growth. In A to C, seedlings were incubated in darkness for 96 h and then illuminated under continuous light for indicated time length. Data are mean ± SE from three biological replicates.

To address the impact of NIP proteins on the RPOTmp function, we grew wild-type and *nip1 nip2* seedlings for 4 d in the dark and illuminated for 2 d under continuous white light and investigated the accumulation of RPOTmp-dependent transcripts in these seedlings (Fig 2B). The patterns of RPOTmp-dependent transcript accumulation were not largely changed between wild type and the double mutant. Because the levels of *trnAUGC* and *trnIGAU* transcripts are also known to be arrested in *rpotmp* mutant (Baba et al. 2004), we also measured the abundance of these tRNA transcripts in the wild-type and *nip1 nip2* seedlings (Fig 2C). The tRNA levels were elevated by light treatment and the patterns were unchanged by the loss of NIP1 and NIP2. These results suggest that the minor influence of NIP proteins in the regulation of RPOTmp functionality during the deetiolaion process.

To test the possibility that NIPs affect the expression levels of RPOTmp, we profiled the mRNA accumulation pattern of *RPOTmp* gene (Fig. 2C). The RPOTmp mRNA levels were increased in the first 6 h of deetiolaion in wild type. The pattern of RPOTmp transcript accumulation was not altered by the mutation of *NIP1* and *NIP2*.

### NIP orthologs are conserved in angiosperms including monocots

Given that RPOTmp is specific to eudicots, we hypothesize that NIPs are also found only in eudicots. To test this hypothesis, we constructed a phylogenic tree of NIP-related polypeptide sequences using those found in a monocot (maize), eudicots (Arabidopsis, spinach, tomato), basal angiosperm (Amborella), gymnosperm (sitka spruce, Japanese cedar), Lycophytes, Pteridophytes, Bryophytes, and green algae (*Klebsormidium nitens* and *Chara braunii*) (Figs 3, S1). Our result demonstrated that NIP-related peptide sequences found in seed plants constitute two clades: one was specific to angiosperms and included Arabidopsis NIP1 and NIP2 and spinach NIP (Clade A), and the other included proteins from gymnosperms and angiosperms (Clade B). Although the bootstrap score of the clade B was marginal, the score of the clade A was high as 99. The clade A is consists of a subclade A1 of eudicot and monocot proteins, a small euducot-specific subclade A2, and a protein from a basal angiosperm (Amborella), indicating that NIP-like proteins are found in eudicots and monocots, and the evolutionary origion of this factor can be traced to the basal angiosperms. Of note, the clade A1 also includes Arabidopsis nuclear-targeted NIP paralog, ABA-RELATED RING-TYPE E3 LIGASE (ARRE) (Wang et al. 2018). Considering that spinach NIP and Arabidopsis NIPs are both plastid proteins, the nuclear localization of ARRE would be a derived feature in this molecular group. We caluculated the percentage of serine and threonine, which are enriched in the transit peptide of plastid-targeted proteins (von Heijne et al. 1989), in the N-terminal sequneces of proteins in clades A and B (Figs 3 and S2). Serine/threonine was relatively poor in N-terminal sequence of ARRE compared to plastid-targeted NIPs in Arabidopsis and spinach. We also found that the N-terminal serine/threonine content was low in one tomato protein and two maize proteins in the subclade A1 and the majority of the proteins in the clades A2 and B, whereas many in the A1 and also the NIP-like protein in Amborella had serine/threonine-rich N-terminal regions, supporting our argument that nuclear-localized proteins in the clade A may have been acquired in the later evolutionary steps.

**Figure 3.**
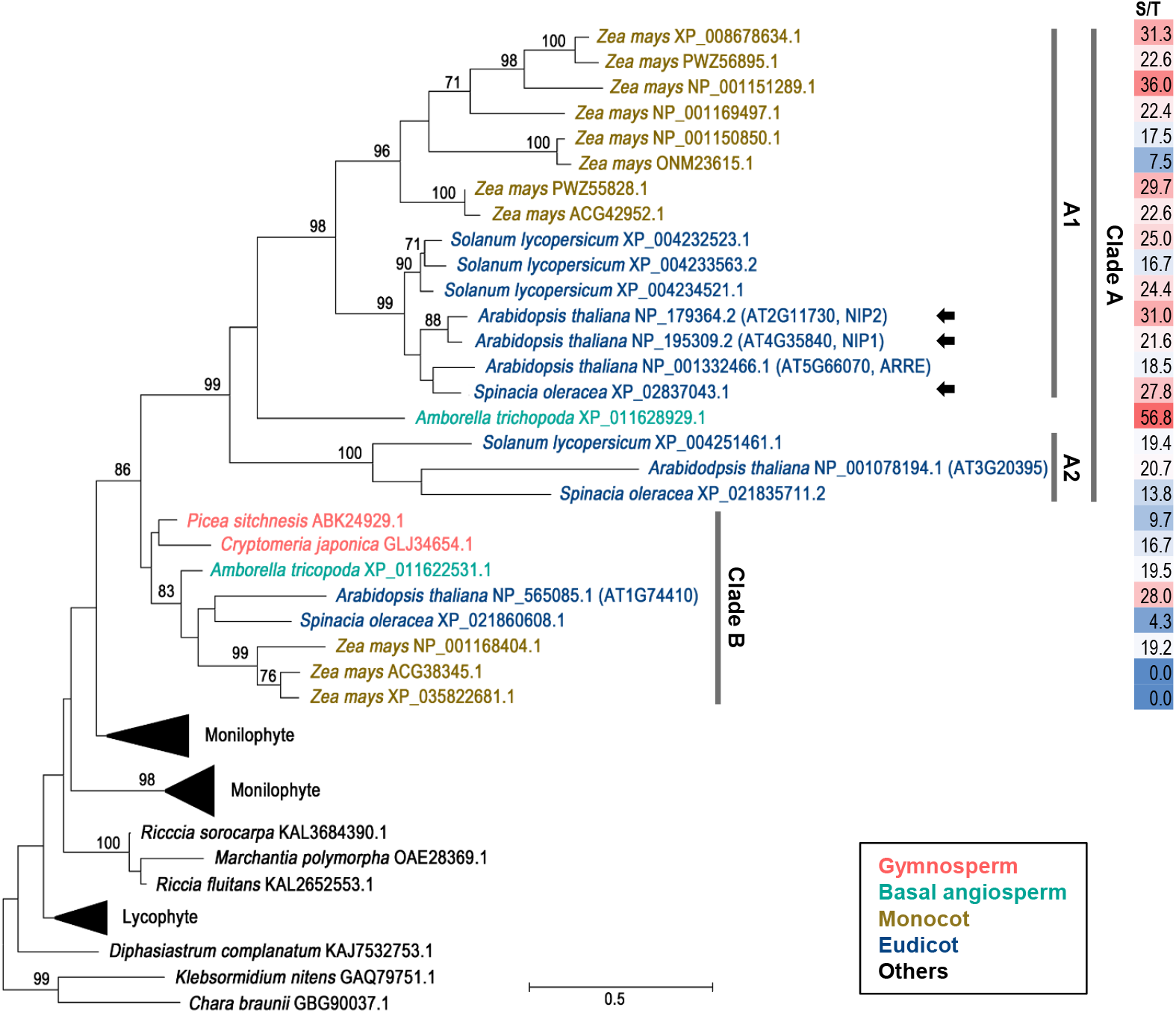
Phylogenetic analysis of NIP. Phylogenetic tree was contracted based on a Maximum Likelihood method with 500 bootstrap replicates. Bootstrap values (≥ 70%) are indicated at each branch. A bar on the bottom indicates distance scale. Black arrows indicate proteins whose plastid localization was experimentally demonstrated. Numbers on the left indicate the percentage of serine/threonine residues in N-terminal regions of proteins in clades A and B. Corresponding regions of the NIP2 transit peptide suggested previously (Azevedo et al. 2008) are used for this analysis. For more details, see supplementary figures S1 and S2.

To estimate the functional redundancy of NIP homologs in Arabidopsis, we aligned the peptide sequences of five NIP-related proteins (Fig. 4A). The overall structure of these proteins including one clade B protein (AT1G74410) may be similar to NIPs, however, their N-terminal region and loop regions between transmembrane domains and the RING domains of AT3G20395 (clade A2) and AT1G74410 (clade B) are distinct from those of NIP1 and NIP2. Alignment score between NIPs and AT3G20395 or AT1G74410 was below 35% (Fig. 4B), indicating different functions between NIPs and these NIP-like factors. Considering the differential localization of NIPs (plastids), and ARRE (nuclei) (Azevedo et al. 2008; Wang et al. 2018), we assume that only NIP1 and NIP2 function in plastids among NIP-related factors in Arabidopsis, although the function and localizaiton of other proteins are remained to be indentified.

**Figure 4.**
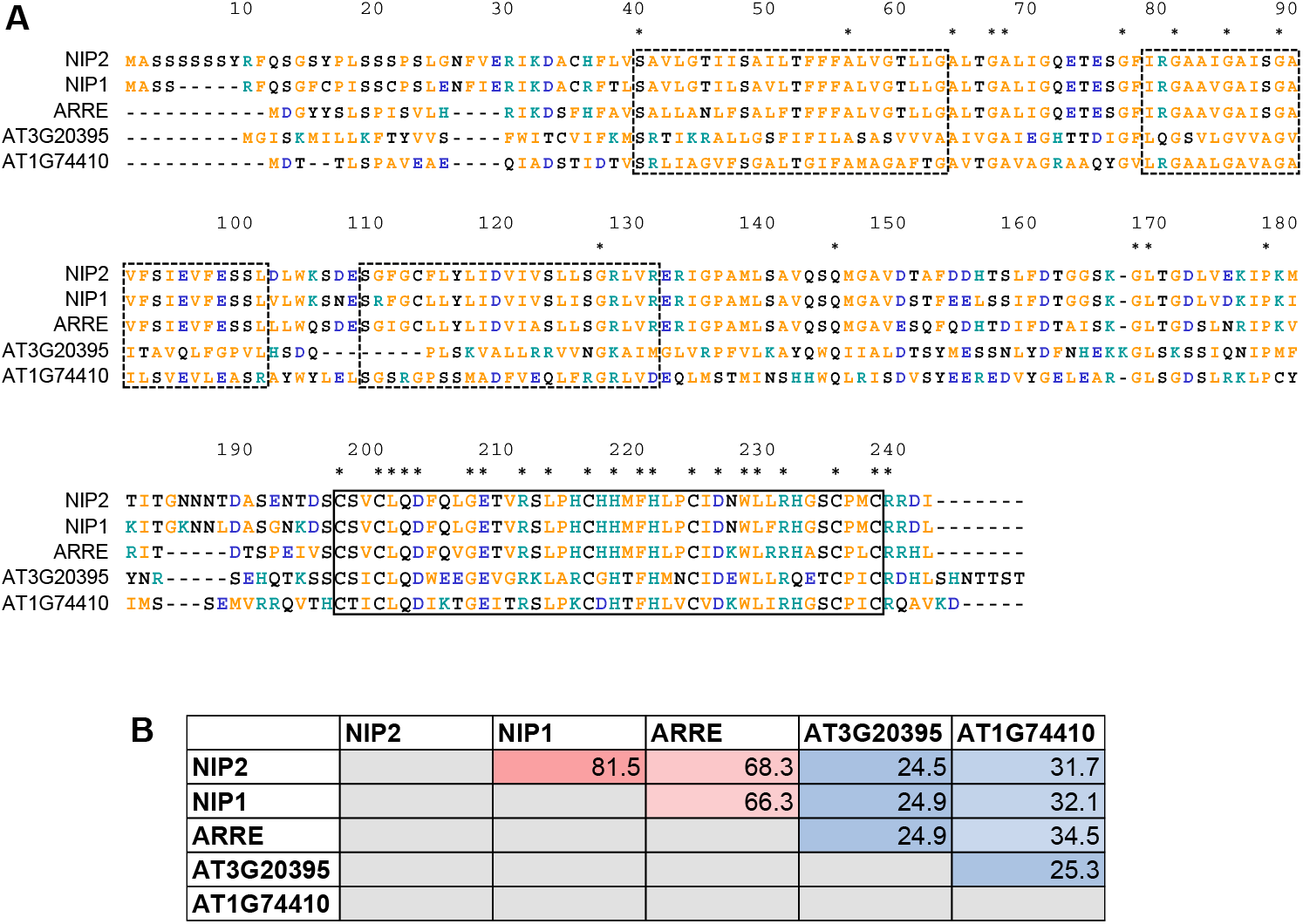
Comparison of five NIP-like proteins in Arabidopsis. (A) Alignment of polypeptide sequences. Positively and negatively charged amino acids are colored in green and blue, respectively. Yellow letters indicate amino acids with hydrophobic side chains, glycine, and proline. Asterisks indicate conserved residues. Boxes with broken and solid lines indicate transmembrane domains and RING domains, respectively, which were reported previously in NIPs (Azevedo et al. 2008). Polypeptide sequences were aligned using CLUSTAL W multiple alignment algorithm mounted in MEGA11. (B) Alignment scores of five NIP-related proteins in Arabidopsis.

## Discussion

### Function of RPOTmp remains stable during etioplast-to-chloroplast differentiation

Consistent with previous studies (Courtois et al. 2007; Azevedo et al. 2008), our RT-qPCR analysis demonstrated that the level of RPOTmp-dependent transcript of *rrn16* was the most abundant in imbibed seeds (Fig 2), suggesting that the role of RPOTmp is prominent in the initiation of plastid differentiation during germination. In contrast to RPOTp which expresses ubiquitously including photosynthetic tissues, RPOTmp is strongly expressed in non-green tissues such as roots and shoot apical meristems (Emanuel et al. 2006), also supporting the idea that RPOTmp plays an important role in immature plastids.

Previous studies showed that the PC promoter activity is decreased within the first 48 h under light (Courtois et al. 2007; Azevedo et al. 2008; Tarasenko et al. 2016). We found a similar decrease of PC-promoter activity in dark-germinated seedlings (Fig 2) thus we conclude that the suppression of RPOTmp during germination is independent of light conditions. We also revealed that the level of *rrn16* transcript was mostly unchanged during the process of etioplast development and etioplast-to-chloroplast differentiation (Fig 2), indicating that the function of RPOTmp remains stable throughout the post-germination stages and is independent of light conditions. Given that chlorophyll accumulation during de-etiolation is retarded by the absence of RPOTmp (Baba et al. 2004) and the activity of PC promoter is consistently higher in the wild type than the *rpotmp* mutant, activity of RPOTmp in etioplasts and developing chloroplasts may also be important for rapid differentiation of chloroplasts after the onset of light illumination.

### Contribution of NIP proteins on RPOTmp regulation is limited during germination and greening

Based on the expression pattern of NIP and the activity of the PC promoter of *rrn16*, Azevedo et al. (2008) proposed a model that induction of NIP expression during chloroplast development suppresses the function of RPOTmp. Our study demonstrated that the mutation of two *NIP* genes in Arabidopsis had no visible impacts on chloroplast development or plant growth (Fig. 1). Moreover, the PC promoter activity was not largely altered during light-triggered chloroplast differentiation in wild type and its pattern was not significantly affected by the loss of NIP1 and NIP2 (Fig 2). From these results, we suggest a modified model of RPOTmp regulation: NIP proteins have limited impact on RPOTmp regulation during chloroplast development, although it may regulate the localization of RPOTmp in plastids and could have some effects on RPOTmp activity in other phases of plastid differentiation.

It may be possible that the *nip1 nip2* double mutant has no visible phenotypes because that additional isoforms of NIP would compensate the regulatory roles of NIP1 and NIP2 in the double mutant. ARRE, the most closely related homolog of NIP, is unlikely to compensate NIPs in plastids, because ARRE was demonstrated to be a nuclear-targeted protein (Wang et al. 2018). Although the function of other NIP-like proteins (AT3G20395 and AT1G74410) in Arabidopsis remains to be addressed, their evolutionary origins and peptide sequences are distinct from those of NIP/ARRE (Figs. 3 and 4), thus we do not expect that these NIP-related proteins function in plastids to regulate RPOTmp. Future studies may identify other RPOTmp regulators and interactors, but they would be structurally and evolutionarily different from these NIP-like proteins.

Regulation of PC promoter activity differs among plant species. In spinach, the PC promoter remains active during etioplast development and is even detectable in chloroplasts, whereas its activity is almost undetectable in tobacco (Iratni et al. 1997). PC-promoter-initiated transcript is also detected in plastids from Arabidopsis leaves and roots (Sriraman et al. 1998), suggesting that Arabidopsis RPOTmp is functional in multiple conditions although the activity may vary and NIPs might be involved in RPOTmp regulation in other developmental stages.

In spinach, another PC-promoter regulator, CHLOROPLAST DNA BINDING FACTOR2 (CDF2) has been reported (Bligny et al. 2000). However, its precise role in transcriptional regulation remains ambiguous (Liere et al. 2011), and its homolog in Arabidopsis has not been reported to date. Compared to RPOTp and RPOTm, RPOTmp has a different amino acid sequence in a promoter-recognizing region (Bohne et al. 2016), suggesting a unique mechanism in its functional regulation. Unidentified features of DNA structures and/or additional regulatory proteins might be involved in the regulation of RPOTmp.

### Evolution of NIP genes may be independent of that of RPOTmp

Phylogenetic analysis of NIP-related polypeptides revealed that angiosperms including monocots, dicots, and basal angiosperms carry proteins structurally similar to well-characterized Arabidopsis and spinach NIPs. We found that the clade for NIP/ARRE (Clade A) is specific to angiosperms and absent in gymnosperms, whereas another NIP-related family (Clade B) was found in gymnosperms and angiosperms (Fig 3). Based on the difference of the evolutionary pattern and peptide sequences between NIPs and clade B proteins (Fig. 4), we assume that clade B factors might have distinct functions from that of NIPs. Alignment scores between an Arabidopsis protein in subclade A2 (AT3G20395) and NIPs were lower than that between a clade B factor (AT1G74410) and NIPs, thus proteins in A2 group also unlikely to have NIP-like functions. ARRE in Arabidopsis have the highest similarity with NIPs in Arabidopsis and spinach, but it is targeted to nucleus but not to plastids (Wang et al. 2018). These arguments were supported by the relatively low serine/threonine percentage in ARRE and proteins in clades A2 and B, compared to those in clade A1 (Fig 3). Moreover, the N-terminal sequence of a clade A protein in Amborella is highly rich in serine and threonine, suggesting that all groups of angiosperms possess NIP-like proteins in their plastids. For these reasons, we consider that plastid-targeted NIP-like proteins were acquired in the common ancestor of angiosperms, whereas Arabidopsis and probably some other angiosperms obtained additional extraplastidic NIP-like proteins in the later evolutionary steps.

RPOTp and RPOTm are conserved in angiosperms including basal angiosperm *Nuphar advena*, whereas RPOTmp is only found in eudicots (Yin et al. 2010; Richter et al. 2014). Taken together with our findings, basal angiosperms and monocots are likely to have plastid-targeted NIP-like proteins without possessing RPOTmp, thus NIP-like proteins in these groups may have functions other than regulating RPOTmp. Considering the structural similarity between RPOTp and RPOTmp, NIP homologs might interact with RPOTp in these plants. It is also possible that NIPs have another function in ancestral angiosperm and became interacting proteins of RPOTmp after the acquisition of RPOTmp in eudicots. In summary, our data suggest that NIP and RPOTmp in angiosperms may have evolved in an independent manner. Future studies will clarify how plants modify the regulatory mechanisms of RPOT proteins in chloroplasts by addressing precise localization of NIP-like proteins in monocots and basal angiosperms and functions of these factors, as well as exploring other interacting partners of phage-type organelle RNA polymerases.

## Materials and methods

### Plant materials and growth conditions

All plants, including *rpotmp* (GABI_286E07) (Kühn et al. 2009), *nip1* (SALK_057350), and *nip2* (SALK_137112), are Columbia ecotype of *Arabidopsis thaliana*. Seeds were surface-sterilized and then vernalized in water for 3-4 days at 4°C in the dark. For dark grown seedlings, plants were incubated for 48 or 96 h in complete darkness after illumination for ∼ 3 h to synchronize the germination. For deetiolated seedlings, plants were incubated under continuous white light (50 μmol photons m^-2^ s^-1^) for indicated time length after 96 h of dark incubation. Seedlings were germinated on half-strength Murashige and Skoog medium with 0.8% (w/v) agar and 1% (w/v) sucrose.

### Genotyping

DNA was extracted from seedlings by boiling at 95°C in buffer containing 50 mM Tris-HCl (pH 8.0), 5 mM EDTA and 0.5 mM KCl. PCR was conducted by using MightyAmp DNA polymerase (Takara, ver. 2) with primers listed in the supplementary table S1.

### Reverse transcription-quantitative PCR (RT-qPCR)

Total RNA was extracted from whole seedlings by using Maxwell RSC Plant RNA Kits (Promega). Reverse transcription was performed by use of ReverTra Ace RT qPCR Master Mix with gDNA remover (Toyobo) and quantitative PCR was conducted by use of Thunderbird Next SYBR qPCR Mix (Toyobo) and Thermal cycler Mx3000P (Agilent) with 200 nM of gene-specific primers (Supplementary Table S2). Amplification and detection were performed in duplicate with following settings: initial denaturation, 95°C, 60 s; [denaturation, 95°C, 5 s; annealing and elongation, 60°C, 30 s] × 40 cycles. Transcript abundance was normalized to the averaged level of *ACTIN8* and *ELOGATION FACTOR 1α* transcripts as the internal control following Pfaffl’s method (Pfaffl 2001).

### Identification of AtNIP orthologous genes

To identify the putative NIP-related protein sequences in Streptophytes, we searched for proteins similar to *Arabidopsis thaliana* NIP2 in *Arabidopsis thaliana*, tomato (*Solanum lycopersicum*), spinach (*Spinacia oleracea*), maize (*Zea mays*), *Amborella trichopoda*, and other non-angiosperm organisms using the BLASTp program. Protein sequences with relatively high similarity were selected based on the alignment scores of > 50% of the transmembrane and RING-domains. Alignment scores among Arabidopsis NIP-related proteins were calculated using CLUSTAL W version 2.1 with default settings.

### Phylogenetic analysis

All the obtained amino acids sequences were aligned using Muscle multiple alignment algorithm mounted in MEGA version 7.0.26. The phylogenetic tree of NIP homologs was constructed with MEGA11 version 11.0.13 using Maximum Likelihood method with 500 bootstrap replications. The phylogenetic tree was visualized with MEGA11 version 11.0.13.

## Acknowledgements

We thank Dr. Michiko Sasabe (Hirosaki University) for critical reading of this manuscript. This work was supported by JSPS Grants-in-Aid for Scientific Research (Nos 22H05075, 24K118134, 25H02302).

## Conflict of Interest statement

The authors declare no conflicts of interest.

## Author contributions

YK designed and performed the experiments, analyzed data, and wrote the manuscript. ASN interpreted the data and complemented the writing. SF conceived the project, design the study, analyzed data, and wrote the manuscript. All authors discussed the results and agreed to the final version of the manuscript.

## Notes

### Competing Interest Statement

The authors have declared no competing interest.

