## Supplementary data for "Membranous interacting partners of phage-type plastid RNA polymerase have limited impact on plastid gene expression during chloroplast development"

**Supplementary Table S1.** Oligonucleotide primers used for genotyping. Primer sets of LP and RP or BP and RP are used to detect target genes without or with T-DNA insertions, respectively.

| Target gene | Primer | Sequence |
|---|---|---|
| GABI_286E07 (*rpotmp*) | LP | ATTGGGCTCCATAAATTCGTC |
| | RP | AGCAGCTCTTTCCCATTCTTC |
| | BP (GABI_o8409) | ATATTGACCATCATACTCATTGC |
| SALK_057350 (*nip1*) | LP | TCACCAAGTTGAAAATCCTGC |
| | RP | ATTCCAATCTGGGTTTTGTCC |
| SALK_137112 (*nip2*) | LP | TTTGCACTGCACTTAGCATTG |
| | RP | CGTTTAAGCACACGACTAGACG |
| SALK_057350<br>SALK_137112 | BP (LBb1.3) | ATTTTGCCGATTTCGGAAC |

**Supplementary Table S2.** Oligonucleotide primers used for quantitative PCR analysis.

| Target gene | AGI code | Sequence<br>Forward | Reverse |
|---|---|---|---|
| *ACT8* | AT1G49240 | ACTGTGCCTATCTACGAGGGTTTC | CCCGTTCTGCTGTTGTGGT |
| *EF1α* | AT1G07940 | TGAGCACGCTCTTCTTGCT | GTGGCATCCATCTTGTTACA |
| *NIP1* | AT4G35840 | GGGCTAAGCAAAACCCATCTC | GGAACCCGCCAAAGATTAAA |
| *NIP2* | AT2G17730 | AAAGCAAACTCACCCCAAGC | CGGTGGGAAGTAGCAGAGTTG |
| $trnA_{UGC}$ | ATCG01190 | TCGGAGAAGGGCAATCACTC | GAGAACCAGGAACGGAGAGC |
| $trnI_{GAU}$ | ATCG01200 | GATGAATCGCTCCCGAAAAG | AGACCTCGCCCGTGAAGTAA |
| *RPOTmp* | AT5G15700 | CAGCATTATGCCGCTCTTGGG | GTCTGCATCTCGGCGCATAATATC |
| *rrn16_PC* | n.a. | GGTAGGGGTAGCTATATTTCTGGGAGC | TGAGCCAGGATCGAACTCTCCATGAG |
| *rrn16_upstream* | n.a. | AGAGGCTCGTGGGATTGACGTGA | TGAGCCAGGATCGAACTCTCCATGAG |

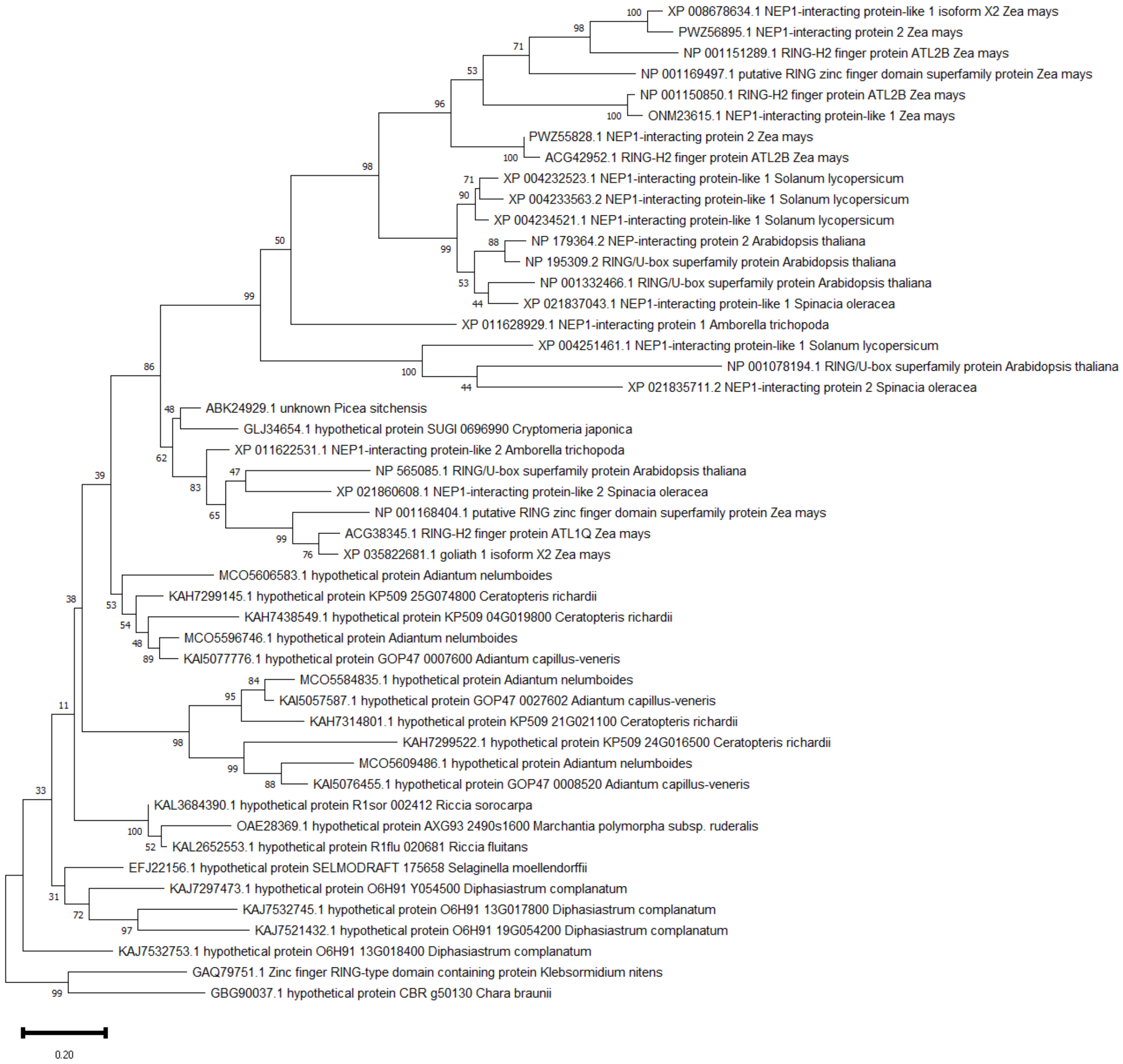


**Supplementary Figure S1**. Phylogenetic tree of all NIP-related polypeptide sequences identified in this study. Bootstrap values are indicated at each branch. A bar on the bottom indicates distance scale.

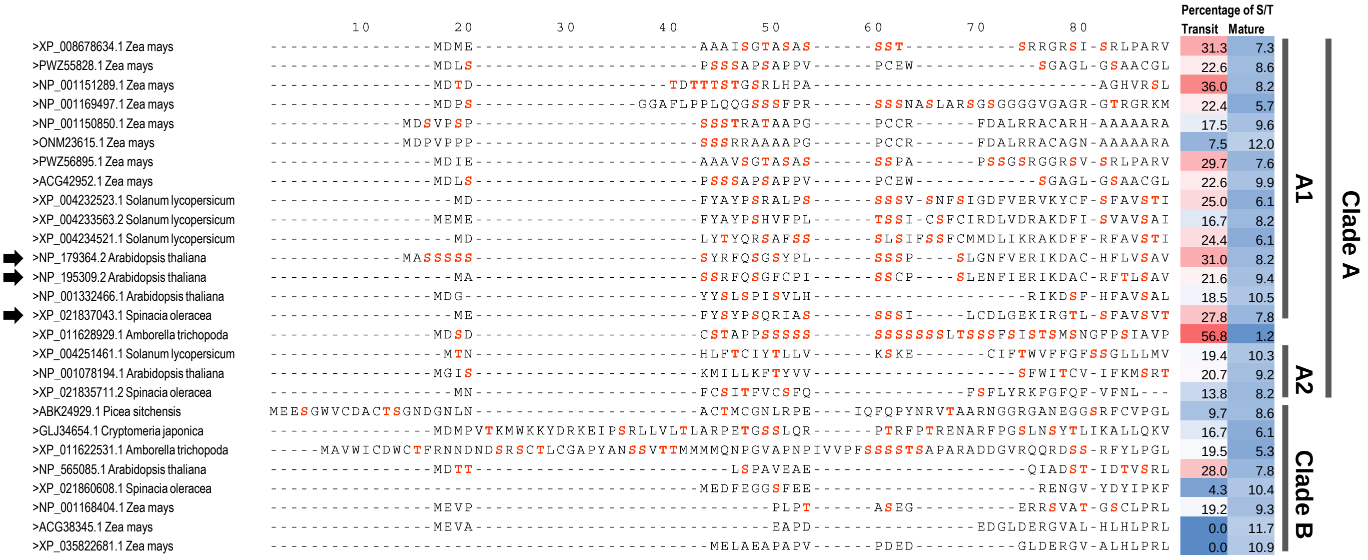


**Supplementary Figure S2**. Alignment and percentage of serine/threonine residues in NIP-like proteins in the clades A and B. Serine/threonine content of N-terminal regions (corresponding regions to the transit peptide of Arabidopsis NIP2/NP_179364.2 and marked as “transit”) and other regions (mature) were calculated. Black arrows indicate proteins whose plastid localization was experimentally demonstrated.
